# Longitudinal profiles of dietary and microbial metabolites in formula- and breastfed infants

**DOI:** 10.1101/2020.05.11.086546

**Authors:** Nina Sillner, Alesia Walker, Marianna Lucio, Tanja V Maier, Monika Bazanella, Michael Rychlik, Dirk Haller, Philippe Schmitt-Kopplin

## Abstract

The early-life metabolome of the intestinal tract is dynamically influenced by colonization of gut microbiota which in turn is affected by nutrition, i.e. breast milk or formula. A detailed examination of fecal metabolites was performed to investigate the effect of probiotics in formula compared to control formula and breast milk within the first months of life in healthy neonates. A broad metabolomics approach was conceptualized to describe fecal polar and semi-polar metabolites affected by diet within the first year of life. Fecal metabolomes were clearly distinct between formula- and breastfed infants, mainly originating from diet and microbial metabolism. Unsaturated fatty acids and human milk oligosaccharides were increased in breastfed, whereas Maillard products were found in feces of formula-fed children. Altered microbial metabolism was represented by bile acids and aromatic amino acid metabolites. Elevated levels of sulfated bile acids were detected in stool samples of breastfed infants, whereas secondary bile acids were increased in formula-fed infants. Co-microbial metabolism was supported by significant correlation between chenodeoxycholic or lithocholic acid and members of Clostridia. Fecal metabolites showed strong inter- and intra-individual behavior with features uniquely present in certain infants and at specific time points. Nevertheless, metabolite profiles converged at the end of the first year, coincided with solid food introduction.

## INTRODUCTION

Nutrition during the early postnatal life is an important factor that might influence health throughout the whole life (Robinson 2015). Breastmilk is the recommended feeding during the first six months, as pointed out by world health organization (WHO n.d.). From a nutritional point of view breastmilk contains an appropriate composition of macronutrients and micronutrients, important for the child’s development. Macronutrients include carbohydrates (lactose, oligosaccharides), fat (triglycerides with saturated and polyunsaturated fatty acids) and proteins (casein or whey proteins such as α-lactalbumin, lactoferrin, secretory IgA, and serum albumin) (Prentice 1996). There is a substantial variation of breastmilk components between mothers depending on diet and age. Infant formula is an industrially produced alternative to breast milk and is used either as a sole food source or to complement breast milk in early life. Infant formulas are based on cow milk, soy or meet special requirements such as hypoallergenic formulas (Koletzko, Baker et al. 2005, Martin, Ling et al. 2016). Feeding of infants with breastmilk or formula is discussed to contribute to different outcomes in health and disease and is a focus in various research fields such as gut microbiome or metabolomics.

A non-targeted metabolomics approach using LC-MS based techniques can be utilized to study diet-related differences between breast- and formula-fed infants. Urine or fecal samples are considered as non-invasive matrices to profile metabolites in health-disease related issues (Wild, Shanmuganathan et al. 2019). Complementary to the gut microbiome research, stool metabolites can reflect changes in gut microbial metabolism. Additionally, excreted host derived metabolites or digested food ingredients provide insights into non-microbial metabolism. In previous work, we are able to show that the fecal metabolome and gut microbiome is altered between breast- and formula-fed infant in the first year of life with converging profiles at the age of 12 months. Probiotic supplementation with bifidobacteria showed only minor effects on both, fecal microbiome and metabolome. Despite the identification of short chain fatty acids, only a shallow description of metabolites was given, with few mass signals and the respective annotated classes (Bazanella, Maier et al. 2017). Here, we extend our non-targeted metabolomics approach by combining different chromatographic approaches to analyze polar and semi-polar metabolites in infant stool and formula or breastmilk, and supporting our metabolite identities with MS based fragmentation experiments and authentic standard matching. The detailed examination between breast- and formula-fed infants allowed us to discriminate between nutrition or microbe derived alterations in the fecal metabolome. Metabolite profiles were monitored at several time points during the first year of life, including a follow-up study one year later, enabling a longitudinal comparison of infant diets.

## MATERIALS AND METHODS

### Study design

Stool samples (n = 244, Table S1) from healthy infants, who received infant formula with (n = 11, F+), without (n = 11, F-) bifidobacteria (*B. bifidum, B. breve, B. infantis, B. longum*) or exclusively breast milk (n = 20, B) were collected over a period of two years in a randomized, double-blinded, placebo-controlled intervention trial as described elsewhere (Bazanella, Maier et al. 2017). Fecal samples of month 1, 3, 5, 7, 9, 12 and 24 were selected for non-targeted analysis. Additionally, breast milk samples (n = 36) were collected at different time points and categorized in samples from secretor (n = 30) and non-secretor mothers (n = 6) (Bazanella, Maier et al. 2017). Furthermore, mother-child and time matching infant fecal samples (n = 31) were selected for comparison of HMO levels between breast milk and associated infant feces. The trial was registered at the German Clinical Trials Register (DRKS00003660) and the protocol was approved by the ethics committee of the medical faculty of the Technical University of Munich (approval number 5324/12).

### Chemicals

Arachidonic acid, eicosapentaenoic acid, myristic acid, 4-hydroxyphenyllactic acid, indolelactic acid, phenyllactic acid, 6’-sialyllactose, chenodeoxycholic acid (CDCA), cholic acid (CA), ursodeoxycholic acid (UDCA), lithocholic acid (LCA), glychochenodeoxycholic acid (GCDCA) and taurochenodeoxycholic acid (TCDCA) were purchased from Sigma-Aldrich (St. Louis, USA). 7-oxoLCA, 3-dehydroCDCA, 7,12-dioxoLCA, 7-oxodeoxycholic acid (7-oxoDCA), 3-dehydroCA, 7-epiCA, glycocholic acid (GCA) and taurocholic acid (TCA) were purchased from Steraloids (Newport, RI, USA). Cholic acid 7-sulfate (CA-S) and 3’-sialyllactose were purchased from Cayman (Biomol GmbH, Hamburg, Germany). Sulfate (S) conjugates of CDCA, UDCA and LCA were synthesized according to Donazzolo et al. (Donazzolo, Gucciardi et al. 2017) and structures were verified by MS/MS and NMR spectroscopy (data not shown). Milli-Q water (18.2 MΩ) was derived from a Milli-Q Integral Water Purifcation System (Billerica, MA, USA). Acetonitrile (ACN; LiChrosolv®, hypergrade for LC-MS), methanol (LiChrosolv®, hypergrade for LC-MS) and ammonium acetate (NH_4_Ac) were obtained from Merck (Darmstadt, Germany). Glacial acetic acid was purchased from Biosolve (Valkenswaard, Netherlands) and formic acid form Honeywell Fluka™ (Morristown, NJ, USA).

### Fecal sample preparation

Metabolite extraction from infant stool samples was prepared with methanol as described previously (Bazanella, Maier et al. 2017). For hydrophilic interaction liquid chromatography (HILIC) analysis the methanol extracts were evaporated under vacuum at 40 °C (SpeedVac Concentrator, Savant SPD121P, ThermoFisher Scientific, Waltham, MA, USA) and reconstituted with ACN/H_2_O 75:25 (v/v). A pooled sample was generated from all fecal extracts for quality control purpose. All samples were stored at −80 °C in tightly closed tubes.

### Breast milk sample preparation

For HILIC measurements, 125 µL breast milk was extracted with 375 µL acetonitrile. The mixture was vortexed, centrifuged with 14,000 rpm at 4 °C for 10 min and the supernatant was collected. A pooled sample was generated from all breast milk extracts for quality control purpose. All samples were stored at −80 °C in tightly closed tubes.

### Quantification of fatty acids in formula and breast milk

Breast milk samples from lactation month 1 (n = 26), month 3 (n = 28) and month 4 (n = 9) were pooled, respectively. Additionally, 3 types of infant formula (pre, 1 and 2), consumed by infants of this study, were mixed with hot tap water (~50 °C) according to manufacturer instructions. The method for quantification of the fatty acids in the different milk samples was described by Firl et al. (Firl, Kienberger et al. 2014).

### UHPLC-MS/MS screening

The fecal and milk extracts and standard substances (in ACN/H_2_O 75:25, v/v for HILIC or methanol for RP) were analyzed by UHPLC (Acquity, Waters, Milford, MA, USA) coupled to a time of flight (TOF) mass spectrometer (MS) (maXis, Bruker Daltonics, Bremen, Germany). HILIC was performed using an iHILIC®-Fusion UHPLC column SS (100×2.1 mm, 1.8 µm, 100 Å, HILICON AB, Umea, Sweden). Chromatographic settings were the same as previously described (Sillner, Walker et al. 2019) with the following modifications: injection volume was 5 µL, eluent A consisted of 5 mmol/L NH_4_Ac (pH 4.6) in 95% ACN (pH 4.6) and eluent B of 25 mmol/L NH_4_Ac (pH 4.6) in 30% ACN with a runtime of 12.1 min, followed by reconditioning for 5 min after each sample, respectively. Every tenth injection a pooled fecal sample was used as quality control for subsequent batch normalization.

Calibration of the MS was done by injecting ESI-L Low Concentration Tuning Mix (Agilent, Santa Clara, CA, USA) prior to the measurements. Additionally, ESI-L Low Concentration Tuning Mix (diluted 1:4 (v/v) with 75% ACN) was injected in the first 0.3 min of each UHPLC-MS/MS run by a switching valve for internal recalibration. Mass spectra were acquired in negative electrospray ionization mode (-ESI), respectively. Parameters of the ESI source were: nitrogen flow rate: 10 L/min, dry heater: 200 °C, nebulizer pressure: 2 bar and capillary voltage: 4000 V. Data were acquired in line and profile mode with an acquisition rate of 5 Hz within a mass range of 50–1500 Da. Data-dependent MS/MS experiments were performed in automated MS/MS mode. After each precursor scan, the five most abundant ions (absolute intensity threshold ≥ 2000 arbitrary units) were subjected to MS/MS. Each fecal sample was measured in duplicates with a collision energy of 10 and 35 eV, respectively. The RP UHPLC-MS measurements for non-targeted metabolomics and short chain fatty acid analysis are described elsewhere (Bazanella, Maier et al. 2017).

Raw UHPLC-MS data were processed with Genedata Expressionist Refiner MS 11.0 (Genedata GmbH, Munich, Germany), including chemical noise subtraction, intensity cutoff filter, calibration, chromatographic peak picking and deisotoping. Metabolite library search and classification was done with the Human Metabolome Database (HMDB) (Wishart, Feunang et al. 2018) for MS1 or MS2 spectra (± 0.005 Da). Manual classification was done for Maillard reaction products.

Peak areas of duplicates were averaged and normalized to fecal weight. Batch normalization based on consecutive quality control measurement samples (pooled sample) was performed after missing value imputation (randomized number between 1.0 and 1.2, based on lowest value of the data matrix).

Targeted MS/MS experiments of Amadori products were performed in multiple reaction monitoring mode (MRM) at 20 eV in positive ionization mode (Sillner, Walker et al. 2019).

### Statistical analysis

Multilevel partial least squares discriminant analysis (PLS-DA) was applied to consider the paired and multi-factorial (diet and time) design of the study. The validity of the multilevel PLS-DA model was confirmed by 7-fold cross validation and receiver operating characteristic (ROC) curves, by evaluating the first and second component. Regularized canonical correlation analysis (rCCA) was applied for correlation between OTU (operational taxonomic unit) and bile acid data (Meng, Kuster et al. 2014). The vertical integration method combined the two datasets (Sperisen, Cominetti et al. 2015). Inter- and intra-individual metabolites were selected by multiple co-inertia analysis, available at the omicade4 package of R environment (Meng, Kuster et al. 2014). Prior all analyses, data was log-transformed and unit-variance scaled.

### High-throughput 16S rRNA gene sequencing

Samples were prepared and analyzed as described previously (Bazanella, Maier et al. 2017).

## RESULTS AND DISCUSSION

### Time independent feeding effects on the fecal metabolome

The particular design of the infant study with longitudinal sampling requires appropriate statistical methods due to high dimensional and co-linear metabolite data. By applying a multilevel PLS-DA analysis, the time dependent effects in this study were eliminated and metabolites representing the effects of the different feeding types, i.e. breastfed (B, blue) or formula-fed (F-/F+, orange or green) were extracted (Fig. 1). The first component of the multilevel PLS-DA explained 4% (HILIC, Fig. 1A) and 3% (RP, Fig. 1B) of the total variance between B and F groups. Despite the clear visual separation of feeding groups, the relatively low percentage of the feeding variable on the total variance can be explained by the fact that infant fecal metabolomes are influenced by many different variables such as the amount of food, gastrointestinal passage or digestion status and sampling time, all contributing to the variability. Probiotic treatment (F+, green) could explain only 1% of the total variance in the metabolite data. Only few infants of the F+ group in month 1 and 3 seems to respond to probiotics, as seen in the right lower corner of Fig 1A and B. Receiver Operator Characteristic (ROC) curves of the multilevel PLS-DA component 1 confirmed with around 0.99 classification accuracy that B are highly distinct from F groups (Fig. S1, A, C). Probiotic supplemented vs. non-supplemented formula feeding, represented in component 2 (Fig. 1), resulted in slightly lower classification accuracy of around 0.98 (Fig. S1, B, D). For further investigations, the most contributing features (top 15%) were selected from the loadings plot, starting from lowest (B) or highest (F - and F+) principle component 1 (p1) coordinates and the same for principle component 2 (p2) coordinates (F- vs. F+) values (Fig. 1C, D).

**Figure 1.**
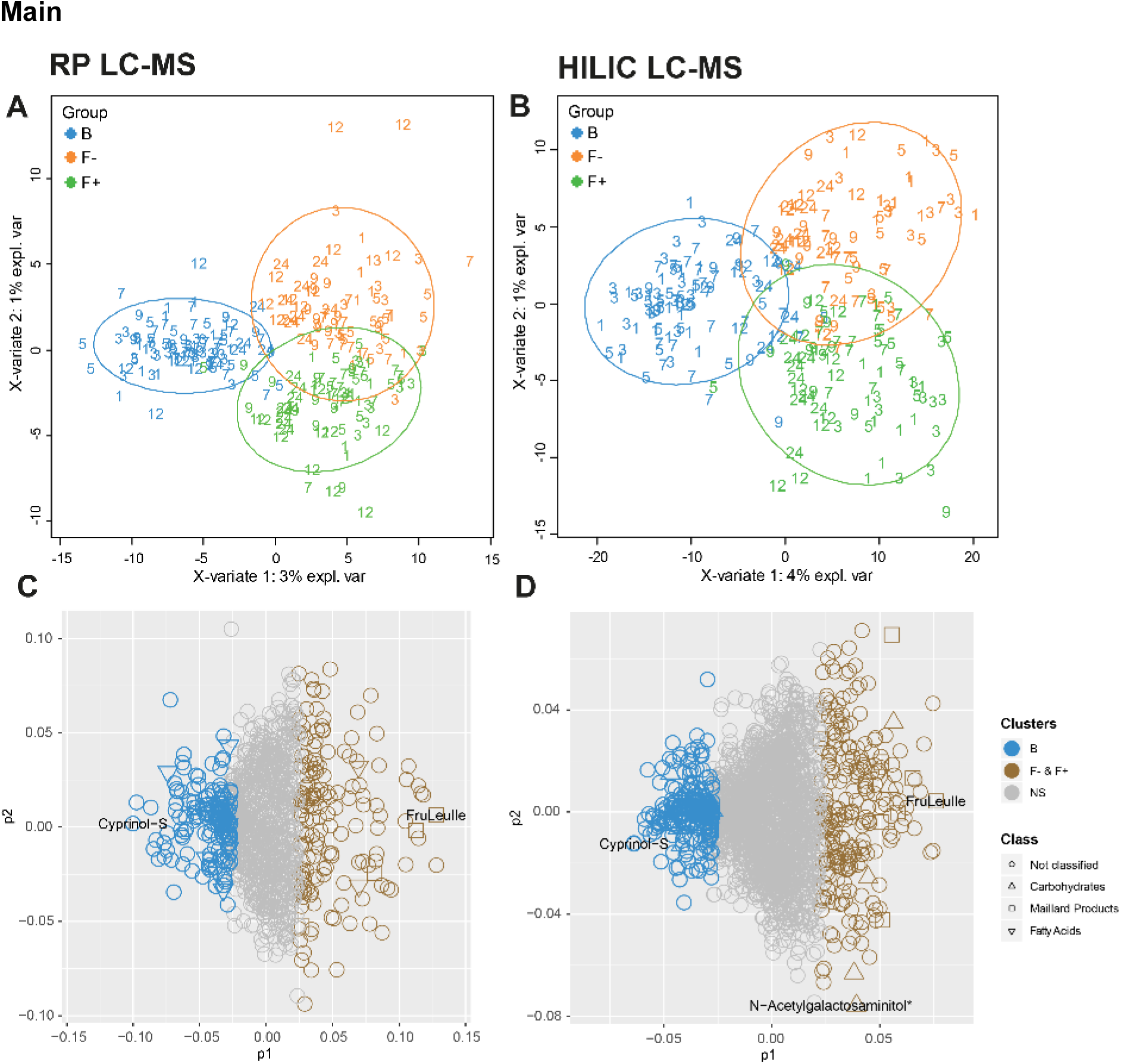
Multilevel PLS-DA. Scatter plots of (A) RP- and (B) HILIC-UHPLC-MS measurements (negative electrospray ionization mode). Groups were separated according to feeding and corrected for time. Numbers represent age of infants at the time of feces sampling (month 1 – 12, 24). Multilevel PLS-DA loading plots for (C) RP and (D) HILIC UHPLC-MS. Top 15% discriminant features were highlighted for B (breastfed, blue) or both F groups (formula-fed, brown). The remaining features are presented in grey (NS=not significant). Putative carbohydrates (open triangle), fatty acids (open inverted triangle) and Maillard products (open box) are representing significant clusters between B and F groups. Cyprinol-S (-sulfate) was increased in breastfed infants, the Amadori product FruLeuIle (*N-*deoxyfructosylleucylisoleucine) was specifically representing both F groups. NAcetylgalactosaminitol was increased in the F+ group. *detected as [M+Cl]^-^ adduct.

Putative annotation of features indicated that group B is described by “*fatty acids and conjugates*” for RP and “*carbohydrates and carbohydrate conjugates”* for HILIC analysis, whereas many characteristic metabolites of both F groups turned out to be Maillard reaction derived compounds, especially Amadori products (Sillner, Walker et al. 2019). During the identification process, features were grouped either in nutrition or microbe derived metabolites, based on existing literature (Bode 2012, Pischetsrieder and Henle 2012, Dodd, Spitzer et al. 2017, Sillner, Walker et al. 2018, Pranger, Corpeleijn et al. 2019). Metabolites were annotated (level 3) or identified (level 1 and 2). The list of all features of the RP and HILIC analysis including the loadings information is available in Tab. S1.

## Nutrition derived metabolites in course of time

### Increased fatty acids in breastfed children reflected fatty acid composition of breastmilk

The semi-polar fecal metabolome of group B was discriminated by the subclass of fatty acids and conjugates. In feces of breastfed infants the saturated fatty acid myristic acid (C14:0) was elevated from month (m) 1 – 5. In formula-fed infants myristic acid was very low abundant during the first 3 months, but showed very similar levels to breastfed infants at month 12 and 24 (Fig. 2A), probably due to the predominant solid food nutrition in all of the groups at this age. Myristic acid was also reported to be higher in cecal contents of breast-fed piglets (Poroyko, Morowitz et al. 2011). On the contrary, a different study reported myristic acid to be significantly higher in formula-fed infants (Chow, Panasevich et al. 2014), which could be due to the variable fatty acid composition of different infant formulas. To verify our findings we quantified the fatty acids in breast milk and infant formula relevant for our study (Tab. S2). Indeed, the average concentration of myristic acid in breast milk was found to be much higher than in in the infant formulas (pre, 1, 2) consumed by our cohort. Furthermore, also the long chain poly-unsaturated fatty acids arachidonic (Fig. 2B) and eicosapentaenoic acid (Fig. 2C) were increased in breastfed infants compared to the two formula-fed groups up to month 7. Again, this relation was confirmed in the fatty acid analysis of breast milk and formula (Tab. S2), showing higher concentrations of arachidonic and eicosapentaenoic acid in breast milk. Although, the total amount of poly-unsaturated fatty acids was higher in formula.

**Figure 2.**
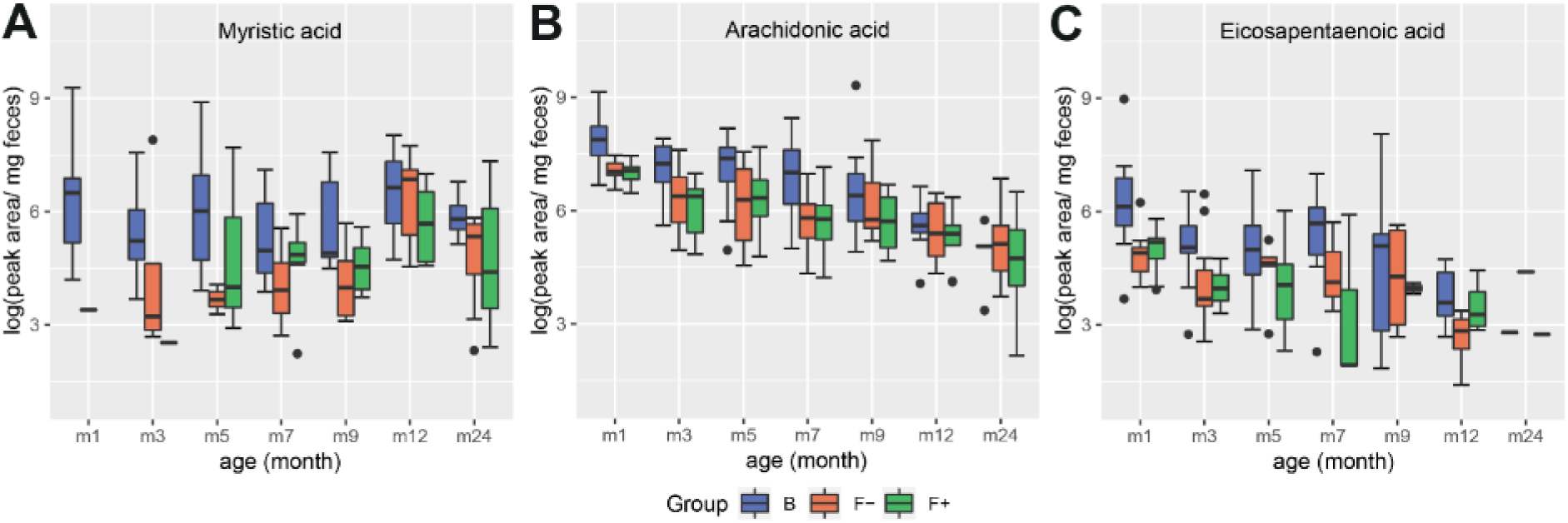
Diet-related alteration of fatty acid profiles in feces of breastfed (B, blue) and formula-fed infants without (F-, orange) or with probiotics (F+, green) over time. (A) Myristic, (B) arachidonic and (C) eicosapentaenoic acid were increased in group B up to month 7.

### Infant fecal HMOs reflected the secretor status of the mothers

In the HILIC analysis, human milk oligosaccharides (HMOs) were very characteristic for the fecal metabolome of breastfed infants. HMOs are only present in human breast milk, therefore these mass signals were absent in the formula-fed group. The HMO profile in breast milk is determined by the secretor status. Non-secretor mothers have an inactive allele of the maternal fucosyltransferase 2 (FUT2) gene and therefore can’t produce α1-2 fucosylated HMOs, e.g. 2’-fucosyllactose. According to this, breast milk samples were classified in secretor (n = 30) and non-secretor (n = 6) samples by Bazanella et al. (Bazanella, Maier et al. 2017). This classification was also done for mother-child matched fecal samples to examine the influence of the mothers secretor status on the HMO profiles in feces of their infants by applying a non-targeted profiling HILIC-MS approach (Sillner, Walker et al. 2019) on both matrices. In total, we detected 6 HMOs (Fig. 3). Unfortunately, no differentiation between 2’-fucosyllactose and 3’-fucosyllactose was possible with the non-targeted screening method, therefore both were summarized as fucosyllactose. The relation of the HMOs in breast milk from secretor vs. non-secretors were very precisely reflected in the fecal samples of infants from the corresponding secretor vs. non-secretor mothers (Fig. 3). Interestingly, this was not only the case for fucosylated but also for sialylated HMOs, like 3’- and 6’-sialyllactose. This illustrates that the mothers secretor status can also be determined by analyzing HMOs in infant stool. The secretor status also influences the infant gut microbiota composition. Feeding with breast milk from secretor mothers enhances the colonization with specific bifidobacteria, which are common infant gut commensals (Lewis, Totten et al. 2015, Bazanella, Maier et al. 2017).

**Figure 3.**
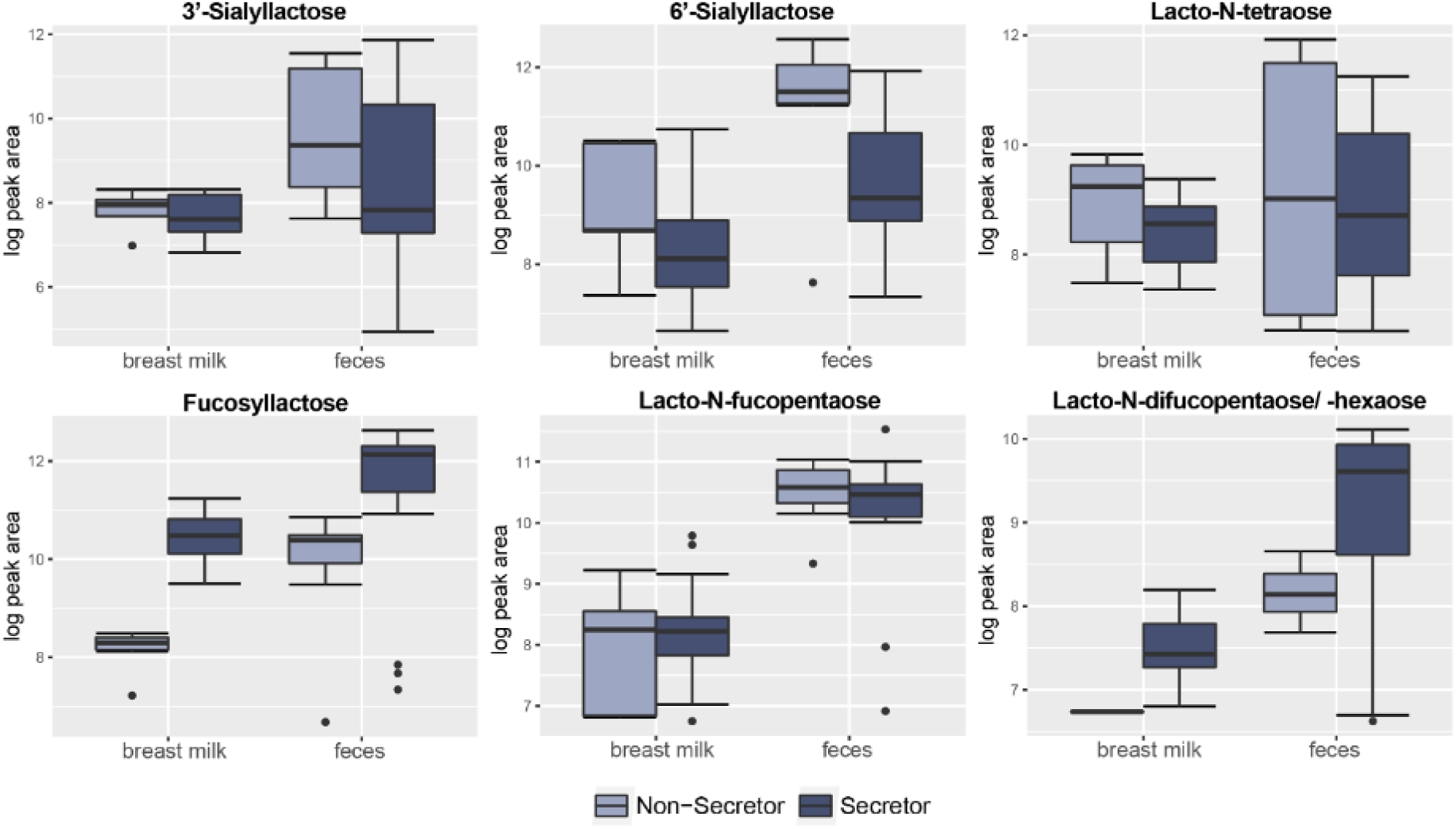
Relation of six different human milk oligosaccharides (HMOs) in breast milk versus feces. Breast milk samples were categorized in secretor (n = 30) and non-secretor (n = 6). Breast milk and feces samples (n = 31) were mother-child and time matched for comparison. The secretor status determined the relative amount of different HMOs in breast milk as well as in the corresponding feces samples.

### Maillard reaction products dominated the stool metabolome of formula-fed infants

Recently, we identified the milk-derived Amadori products *N*-deoxylactulosyl- and *N-*deoxyfructosyllysine and *N*-deoxylactulosyl- and *N*-deoxyfructosylleucylisoleucine in stool from the same cohort, which were only present in formula-fed infants (Sillner, Walker et al. 2019). Amadori products are early Maillard reaction products and are formed during infant formula production, mainly between lactose and protein bound amino acids (N-terminal or lysine side chains). We were able to annotate further Amadori products as significant features of metabolome the formula-fed children, due to their characteristic MS/MS patterns (Fig. 4). The *m/z* signal 381.1338 was only present in formula-fed infants and decreased slowly with time, most likely due to solid food introduction (Fig. 4A, Tab. S1). MRM fragmentation in positive mode of the corresponding *m/z* signal 383.1481 is shown in Fig. 4B. Dominant fragment ions were neutral losses of water, loss of 84 Da (−3H_2_O-CH_2_O) and loss of the glucose moiety (162 Da), which are characteristic for the Amadori compound class (Hegele, Buetler et al. 2008, Wang, Lu et al. 2008, Ruan 2018, Sillner, Walker et al. 2019). Accordingly, this metabolite was annotated as *N-*deoxyfructosylmethionylalanine (FruMetAla) due to its exact mass (± 0.005 Da mass tolerance), MS/MS fragmentation pattern and the matching N-terminal amino acid sequence MetAla of the formula ingredient glycomacropeptide (Neelima, Sharma et al. 2013). Glycomacropeptide is released from κ-casein during whey powder production and remains in the sweet whey fraction, which is among others used for infant formula (Rigo, Boehm et al. 2001). Furthermore, two peaks were annotated as *N-*deoxyfructosylacetyllysine (FruAcLys) (Fig. S2). Their fragmentation patterns are similar, however the earlier eluting peak (6.2 min) exhibits the typical loss for Amadori products of 84 Da (Fig. S2A), whereas the peak at 6.6 min shares also fragments with *N-*deoxyfructosyllysine (FruLys) (Fig. S2B) (Sillner, Walker et al. 2019). During the Maillard reaction 1-deoxy-2,3-hexodiulose reacts with lysine and degrades to acetyllysine via β-dicarbonyl cleavage (Smuda, Voigt et al. 2010, Henning, Smuda et al. 2011). Since lysine has two amine groups, formation of an acetyllysine Amadori product could be possible during infant formula production. Indeed, we were able to detect peaks with the same mass and retention time in a model reaction mixture of L-lysine with glucose heated for 1 h at 100 °C in water, maybe deriving from glycated α- and ε-acetyllysine (Fig. S2C). FruAcLys and especially FruMetAla could serve as specific nutrition markers for formula-fed infants, similar to the previously proposed *N*-deoxyfructosyl-/*N-*deoxylactulosylleucylisoleucine (Sillner, Walker et al. 2019).

**Figure 4.**
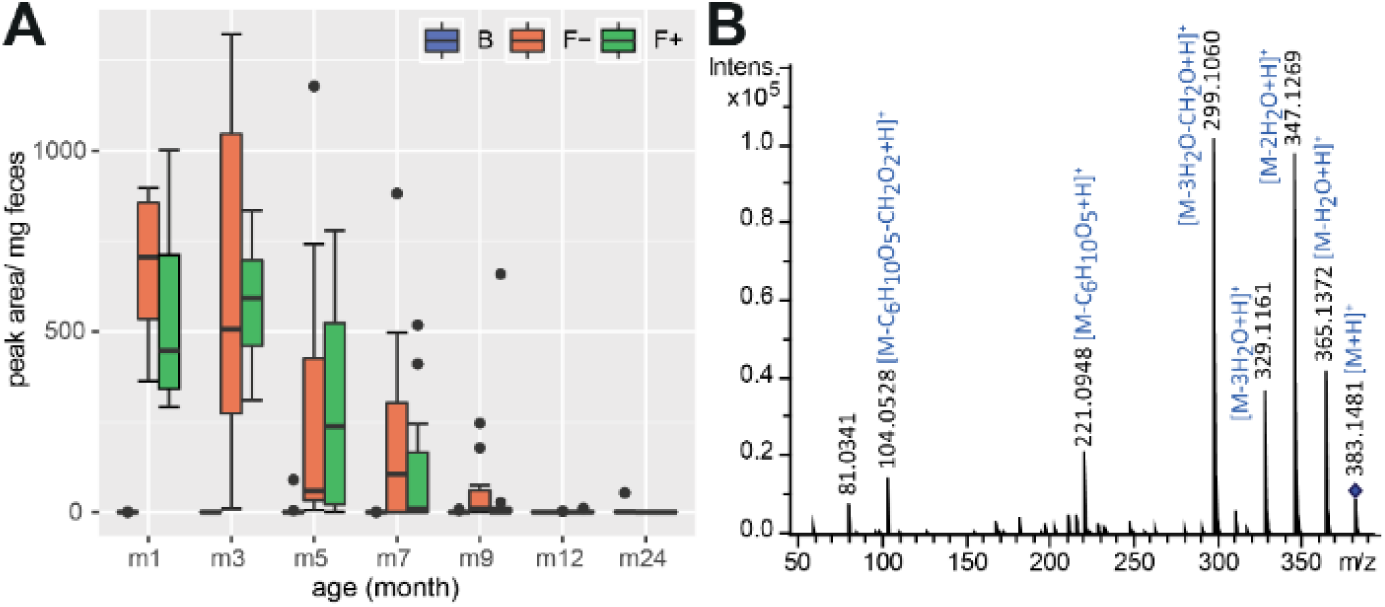
(A) Fecal excretion profile of the putative Amadori product FruMetAla during the first 2 years of life. In formula-fed infants (F-, without probiotics, orange and F+, with probiotics, green) the amount of excreted FruMetAla decreased over time. In breast-fed infants (B, blue) FruMetAla was not detectable. (B) Collision induced dissociation MS/MS experiment (20 eV, positive ionization mode) of FruMetAla in a pooled fecal sample.

## Microbial derived metabolites in course of time

### Diet and time dependent alteration of microbial aromatic amino acid metabolites

During the first year, 4-hydroxyphenyllactic and indolelactic acid (Fig. 5A, B) were increased in fecal samples of breastfed infants. Although 4-hydroxyphenyllactic and indolelactic acid were not significantly changed between F- and F+, higher intensity levels were observed for the bifidobacteria-supplemented F+ group at month 3 and 5. It was reported that bifidobacteria, which are predominant gut microbes in breastfed children, produce phenyllactic, 4-hydroxyphenyllactic (Beloborodova, Khodakova et al. 2009, Beloborodova, Bairamov et al. 2012) and indolelactic acid (Aragozzini, Ferrari et al. 1979) from the aromatic amino acids phenylalanine, tyrosine and tryptophan, respectively. However, the profiles of the corresponding amino acids didn’t show any coherent behavior in feces (Fig. S3), probably due to fluxes into multiple metabolite pathways. Interestingly, phenyllactic acid showed a very different profile compared to the others with no significant differences between the feeding groups but increasing levels over time (Fig. 5C). This may indicate that other bacterial species are settling the children’s gut in the course of time, which are able to produce higher amounts of phenyllactic acid. Whereas, 4-hydroxyphenyllactic and indolelactic acid almost disappeared completely during solely solid food nutrition (24m).

**Figure 5.**
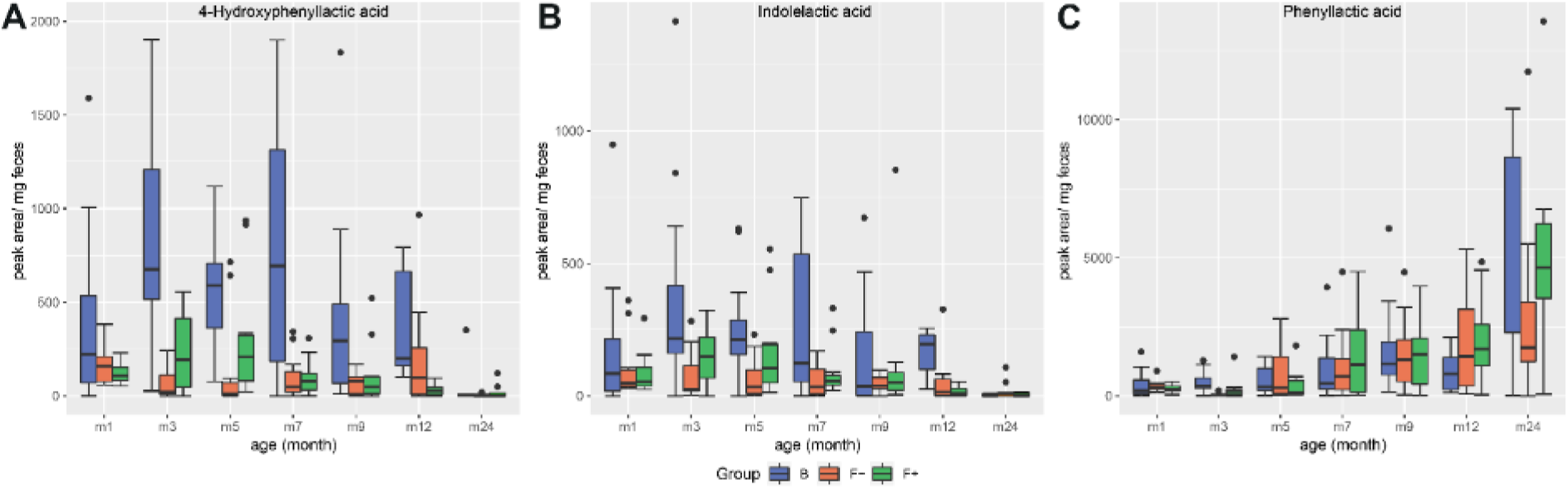
Profiles of bacterial aromatic amino acid degradation metabolites in feces of breastfed (group B, blue) and formula-fed infants without (group F-, orange) or with probiotics (group F+, green) over time. (A) 4-Hydroxyphenyllactic and (B) indolelactic acid were increased in group B up to month 7. (C) Fecal phenyllactic acid increased over time.

### Formula feeding increased the variety of microbial secondary bile acids

Furthermore, several bile acids were significantly affected by the different feeding types. All detected bile acids and the bile acid precursor cyprinol sulfate (cyprinol-S) are displayed in dependence of feeding and age in Fig. 6. Interestingly, the common secondary bile acid LCA was almost absent until month 12. It was already described that the concentration of secondary bile acids are much lower in children compared to adults (Huang, Rodriguez et al. 1976) and that LCA producing bacteria (7α-dehydroxylation) are at first established in the gut at the age of 12 – 18 months (Eyssen 1973). Hammons et al. were able to detect LCA in some infants already at the age of approximately 3 months but with a high inter-individual variation (Hammons, Jordan et al. 1988).

**Figure 6.**
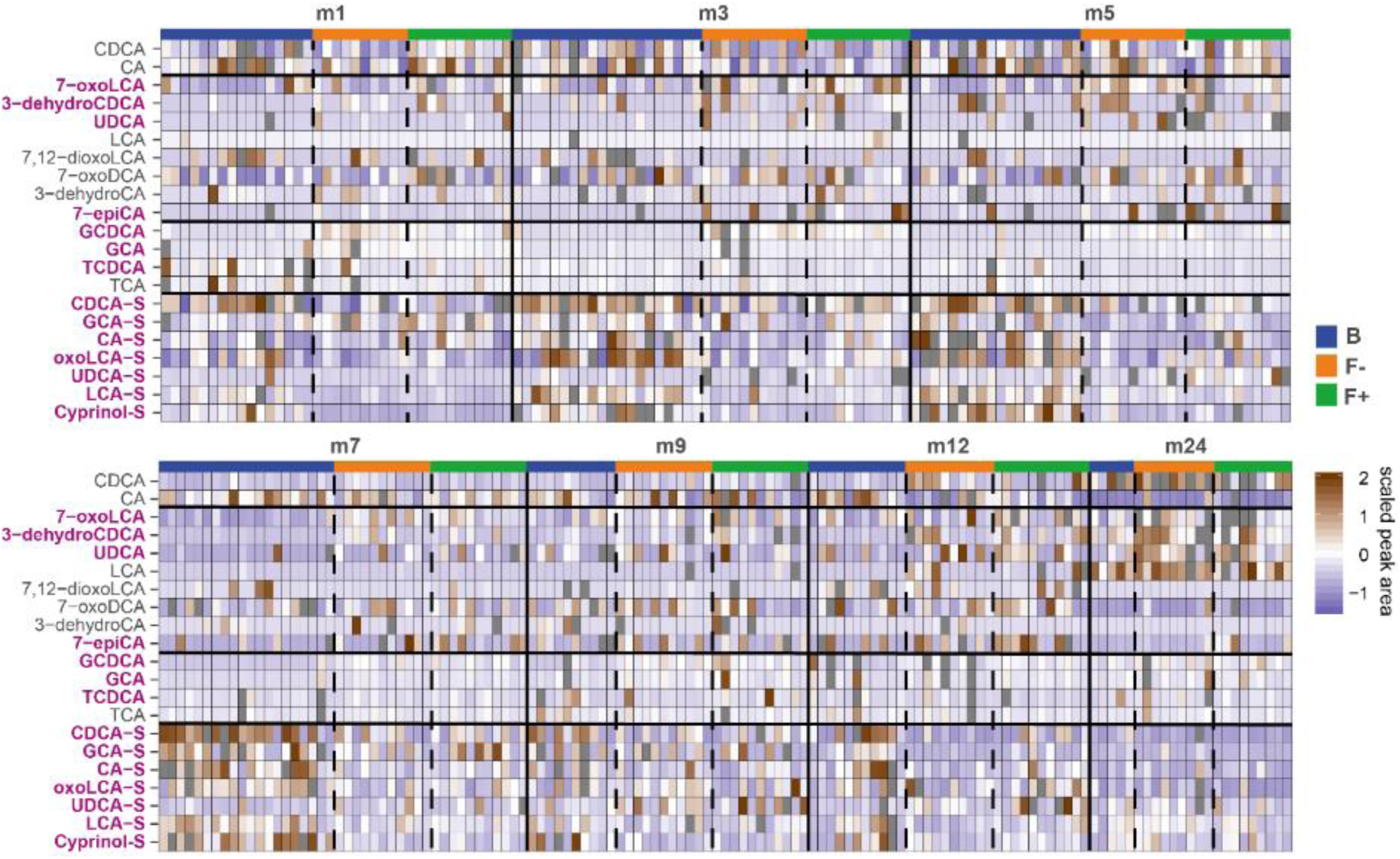
Heatmap of all detected bile acids in feces of breastfed (B, blue) and formula-fed infants without (F-, orange) or with probiotics (F+, green) over time. Peak areas (RP-UHPLCMS) were unit variance scaled. Scaled peak areas > 2 were removed and displayed grey for better visualization. Bile acids which were altered due to the different diets (from multilevel PLS-DA) are marked purple.

Instead, other secondary bile acids were detected in the earlier months, namely 7-oxoLCA, 3-dehydroCDCA, UDCA, 7-12-dioxoLCA, 7-oxoDCA, 3-dehydroCA and 7-epiCA. Lester et al. already reported in the past about the presence of “unconventional” bile acids in infant feces, like UDCA (Lester, St Pyrek et al. 1983). However, 7-oxoLCA, 3-dehydroCDCA, UDCA and 7-epiCA were higher in the formula-fed groups (F+ and F-). This could be due to the fact, that formula-fed children showed a higher bacterial richness and diversity compared to breastfed (B) (Bazanella, Maier et al. 2017). A higher excretion of secondary bile acids in formula- compared to breastfed infants was also reported by Hammons et al. (Hammons, Jordan et al. 1988). Group B, however, was strongly characterized by sulfated bile acids during the first year. Sulfation of bile acids is a major detoxification pathway in humans, which increases their solubility, decreases intestinal absorption, and enhances fecal and urinary excretion. In human adults, it was reported that more than 70% of bile acids in urine are sulfated, whereas the amount in feces is much lower (Alnouti 2009). The high amount of sulfated bile acids in feces of breastfed infants could be due to a defense mechanism against bile acid accumulation, which is suggested in cholestatic diseases (Alnouti 2009), and a decreased absorption rate in the large intestine compared to adults. Heubi et al. reported normal enterohepatic circulation of bile acids to begin at first at the age of 3 – 7 months (Heubi, Balistreri et al. 1982). The conjugated primary bile acids GCA and GCDCA were increased in F- during month 1 and 3. Bifidobacteria show bile salt hydrolase (BSH) activity and are able to deconjugate bile acids (Grill, Manginot-Dürr et al. 1995, Tanaka, Hashiba et al. 2000, Kim, Miyamoto et al. 2004, Begley, Hill et al. 2006, Degirolamo, Rainaldi et al. 2014). The intensities of GCA and GCDCA in the probiotic F+ group during month 1 and 3 were lower and more similar to group B, which could be a sign of probiotic activity.

Regularized canonical correlation analysis (rCCA) (González, Déjean et al. 2008, González, Cao et al. 2012) was performed to explore relationships between bile acids and OTU data (Fig. 7A). A representation of variables defined by the first two canonical variates is displayed in Fig. S4. On the basis of the clustered image map of the cross-correlation matrix (Fig. 7B) and relevance networks were generated (Fig. 7C). Correlations between OTUs and bile acids with a strength > ± 0.3 were found for CDCA, LCA and 3-dehydroCDCA. Correlation strengths > ± 0.5 were only calculated for CDCA and LCA (Fig. 7B). Solely positive correlations were found within these limits. Most of the correlating OTUs belong to the families *Lachnospiraceae* and *Ruminococcaceae*, both of them belong to the class of Clostridia. It is known that many Clostridia species are able to produce the secondary bile acid LCA from the primary bile acid CDCA via 7α-dehydroxylation. The secondary bile acid 3-dehydroCDCA is also produced from CDCA but via 3α-hydroxysteroid dehydrogenases, which were detected in Firmicutes (Fiorucci and Distrutti 2015).

**Figure 7.**
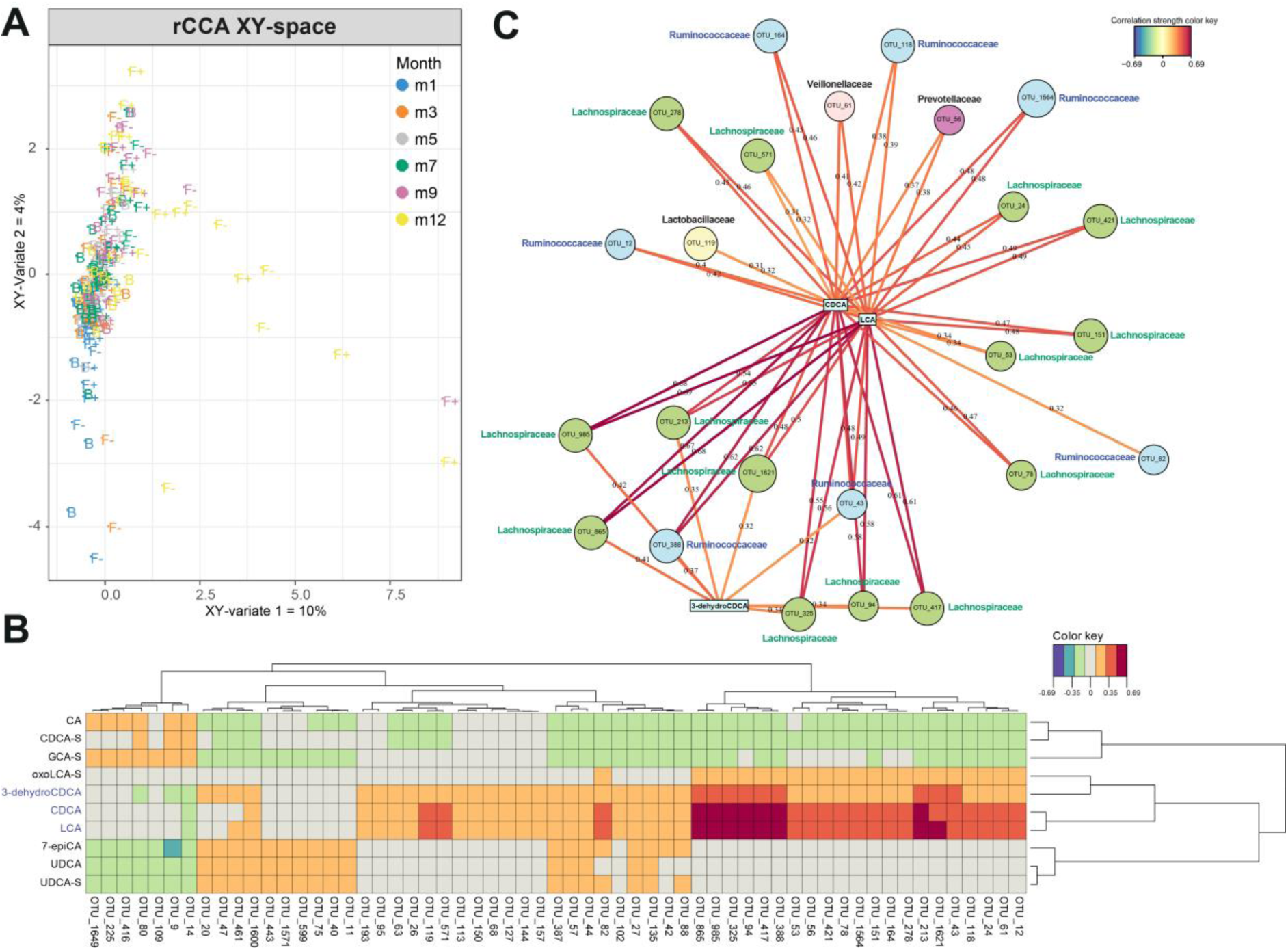
Regularized canonical correlation analysis (rCCA) was used to investigate the correlation between two datasets including LC-MS peak area based bile acid data (X) and OTU data from 16S rRNA sequencing (Y). (A) XY-sample plot is colored according the months and labeled according to the feeding type.(B) Clustered image map of the cross-correlation matrix revealed a distinct cluster of positively correlating OTUs with LCA, CDCA and 3-dehydroCDCA. (C) Corresponding relevance networks for correlations between the two datasets (XY). OTUs are represented by round and metabolites by rectangular nodes. Edges are correlation values, derived from rCCA. Correlation strength > ± 0.3 represented by edges was found for LCA, CDCA and 3-dehydroCDCA. Solely positive correlations were found within these limits. Most of the correlating OTUs belong to the families *Lachnospiraceae* (green nodes) and *Ruminococcaceae* (blue nodes), both of them belong to the class of Clostridia.

### Distinct time and diet inter-and intra-individual metabolite profiles

Longitudinal studies in microbiome research often described highly inter- and intra-individual variability (Zhou, Sailani et al. 2019). The metabolite profiling of stool samples over time enables to determine the variability of each individual and metabolites that are responsible for this behavior. The inter- and intra-individual behavior of the children was elaborated by a multiple co-inertia analysis for the polar metabolome (Fig. 8) (Meng, Kuster et al. 2014). For the exclusively breastfed group (B) only months 1 – 9 were taken into account because of a reduced number of available samples in later months due to weaning. In the breastfed group (Fig. 8A, Tab. S3), especially infant 66 showed very individual profiles at certain time points. Metabolites responsible for this behavior were for example 3-hydroxyphenylacetic acid in month 5, taurocholic acid in month 7 (Fig. S6A) and N-acetylneuraminic acid in month 9. Infant 126 and 115 were characterized by fucosyllactose in month 3 and ferulic acid sulfate in month 7.

**Figure 8.**
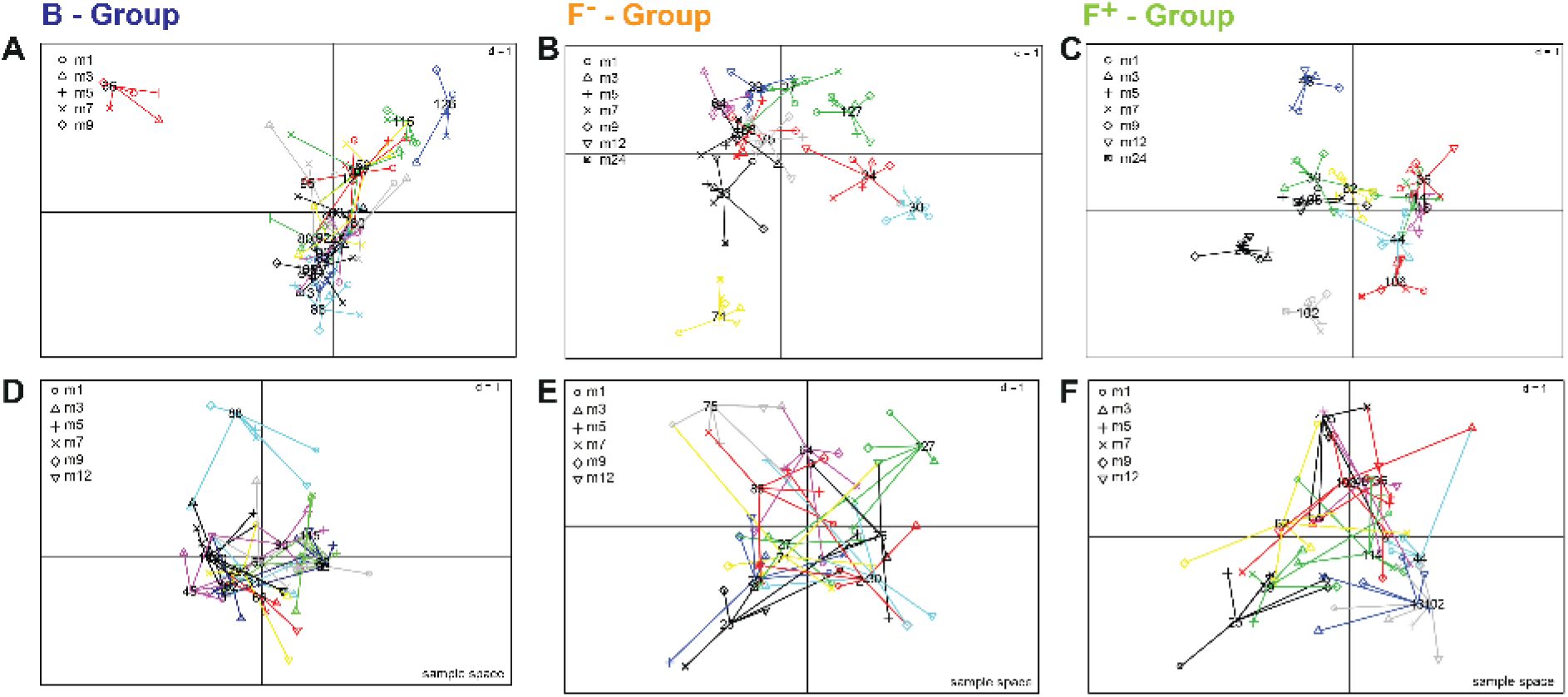
Sample spaces of multiple co-inertia analysis to visualize inter- and intra-individual differences over time, shown for the whole fecal polar metabolome screened with HILIC LCMS approach (A-C) and for short chain fatty acids as selected microbial metabolites (D-F). Each individual (number) is projected within its corresponding feeding group; (A, D) breastfed, (B, E) formula-fed without probiotics and (C, F) formula-fed with probiotics. For the exclusively breastfed group (A) only months 1 – 9 were taken into account because of a reduced number of available samples due to weaning. Samples from the same individual are linked by edges and the shapes represent the different time points. The shorter the edge, the higher the similarity of samples from the same individual. The more similar the metabolic profiles of individuals were the closer the projection in the sample space.

In the F- group (Fig. 8B, Tab. S3) the individuality was less distinct. For infant 71 hydroxydecanoic acid (Fig. S6B) and polyphenolic compounds can be found in month 9 and 12, respectively. The individual profile of infant 30 is characterized e.g. by pantothenol in month 3, which is often used in care products, and fucose in month 9.

In the F+ group (Fig. 8C, Tab. S3) infant 43 showed a different behavior over time due to e.g. cyclamic acid (Fig. S6C), an artificial sweetener in month 3 and 3-hydroxyphenylacetic acid in month 9, whereas infant 25 and 102 were characterized by cyclamic acid in month 12 and 3-(4-hydroxyphenyl)-propionic acid in month 24.

The more differentiated positions of the metabolites in the loading plot of group B and F+ (Fig. S5A, C) compared to the F- group (Fig. S5B) indicates a more pronounced individuality in group B and F+. For group B this could be explained by the fact that breast milk composition can be quite variable between mothers and over lactation periods (Villaseñor, Garcia-Perez et al. 2014, Hascoët, Chauvin et al. 2019, Hewelt-Belka, Garwolińska et al. 2019, John, Sun et al. 2019), whereas the formula-fed infants always received the same type of formula during the trial. Besides xenobiotics like cyclamic acid, the individual behavior of the F+ group seemed to be more influenced by microbial metabolites. Wandro et al. reported that the inter-individual effect on the metabolome of infants over the first 6 weeks of life was stronger than any trends in clinical factors. Furthermore, for some individual infants strong shifts in the metabolite profiles were observed over time, while others remained more stable (Wandro, Osborne et al. 2018). Not only the whole metabolome showed high intra- and inter-individual variability but also specific metabolite classes such short chain fatty acids (McOrist, Miller et al. 2011). Infant 88 (B, Fig. 8D), 75 (F-, Fig. 8E) and 25 (F+, Fig. 8F) showed highest inter- and intra-individual behavior of SCFAs. High dispersion of lactic acid over time was observed in the loading plot of B group (Figure S7, A), but also for F- and F+ infants (Figure S7, B-C). Butyric and propionic acid were also highly distributed over different time points in F- an F+ infants. Valeric, isovaleric and pyruvic showed a more closed distribution for all feeding groups.

In summary, the fecal metabolite profiles of the infant cohort was largely composed of ingredients of consumed food and microbial metabolism of food compounds or co-microbial metabolism of host derived metabolites (Fig. 9). Some of them are recovered directly without modification in stool such as long chain fatty acids or HMOs and may be directly linked to the child’s or mother’s diet or genotype. Furthermore, digested products of food processing (glycated proteins) were found especially in feces of formula-fed infants.

**Figure 9.**
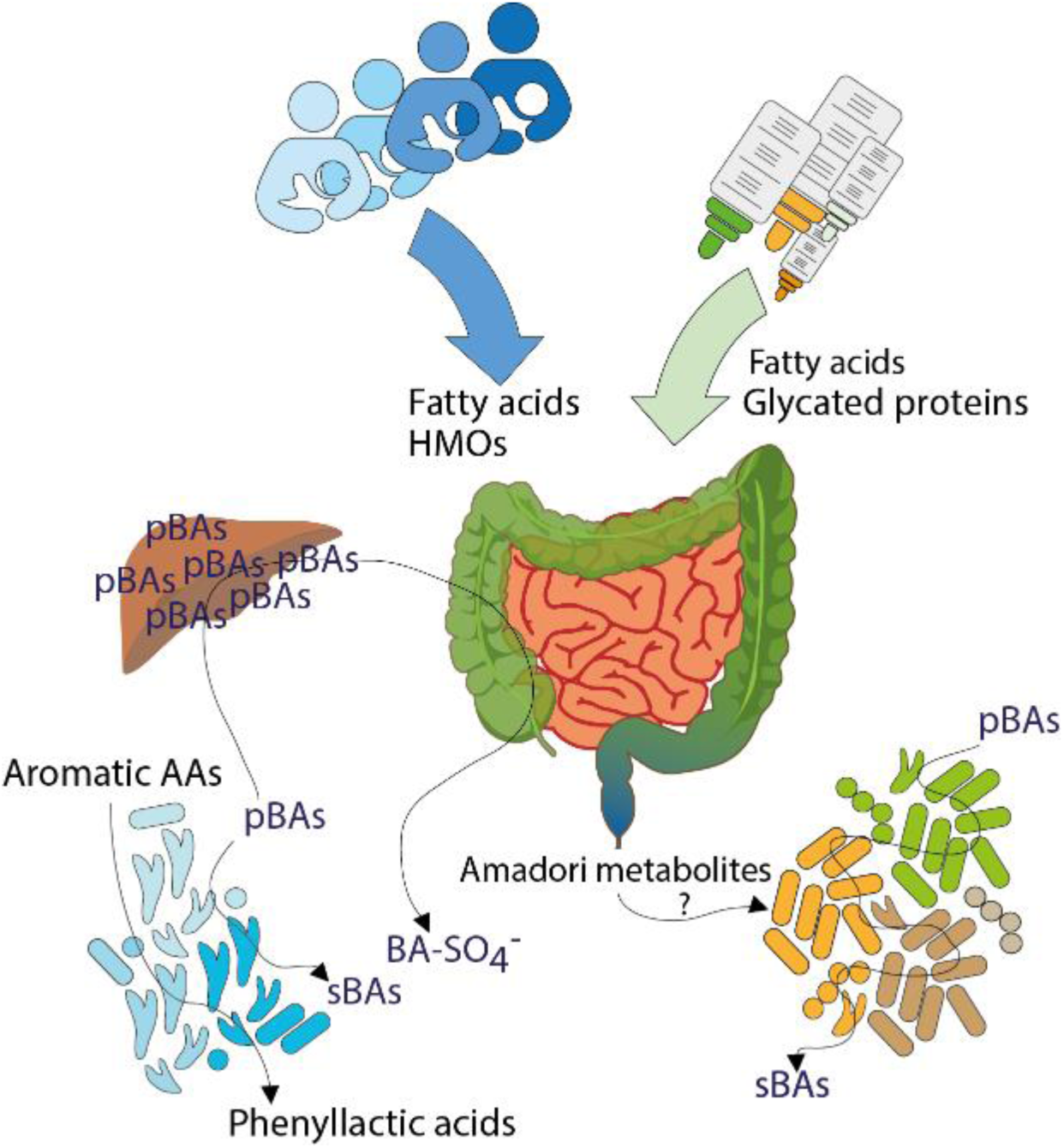
Influences of early life nutrition on the fecal metabolome of infants. Dietary ingredients, endogenous compounds but also microbial products are shaping the gut environment of breast- and formula-fed infants. (pBAs = primary bile acids, sBAs = secondary bile acids, BA-SO_4_^-^ = bile acid sulfates, AAs = amino acids)

Altered microbial modification of amino acids or bile acids in breastfed and formula-fed infants resulted in distinct metabolite profiles, hinting towards different gut microbial populations and thus different microbial metabolism. It may reflect rather specialized or absent microbial metabolism for example of aromatic amino acids through bifidobacteria or increased sulfation of bile acids in breastfed children, respectively. Diverse microbial metabolism was illustrated by several secondary bile acids in formula-fed children. Overall, each individual was reflected by his own fecal metabolite profile and particular metabolites displayed high inter- and individual variability, induced by different genetic backgrounds or external influences, like the individual composition of breastmilk, the amount of consumed formula or living environment.

## Supplementary Information

### Supplementary figures

**Figure S1.**
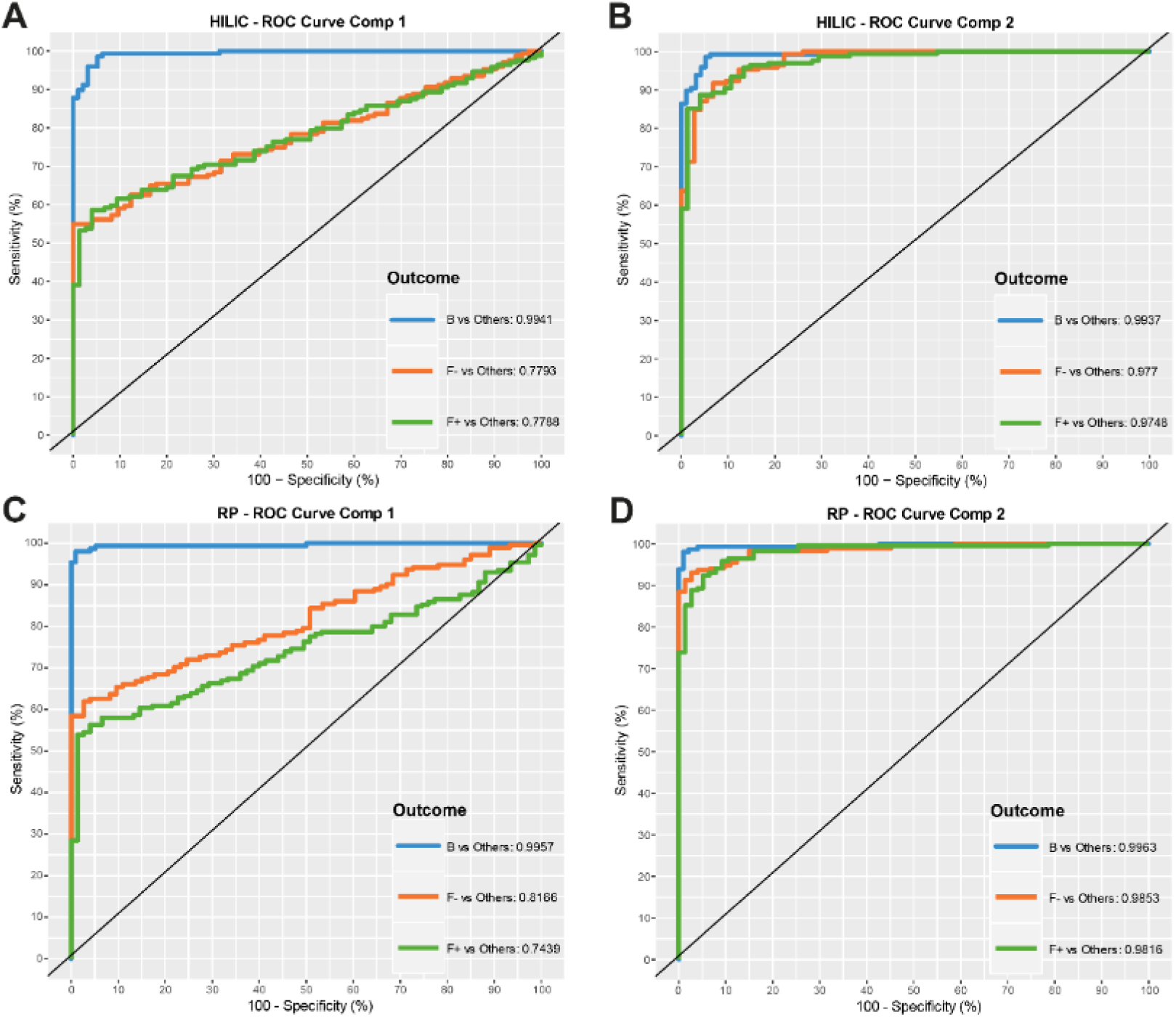
ROC (Receiver Operating Characteristics) curves from the multilevel PLS-DA of (A) HILIC principle component 1, (B) HILIC principle component 2, (C) RP principle component 1 and (D) RP principle component 2 UHPLC-MS measurements (negative ionization mode). The area under the curves indicate a very good separation of breastfed (B) versus formula-fed (F- and F+) infants for both data sets in principle component 1 and 2. Less valid differences between F- and F+ were found in component 2.

**Figure S2.**
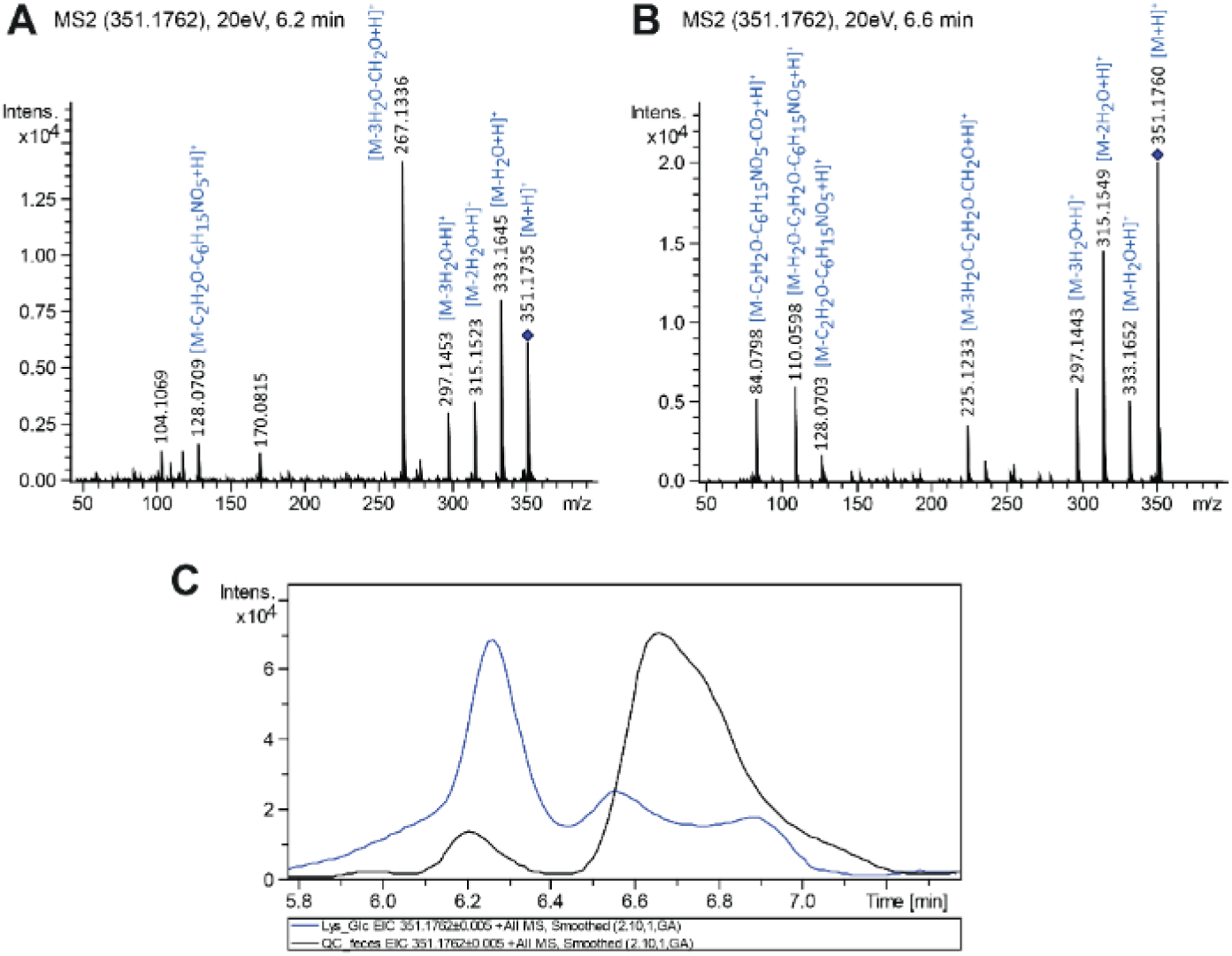
Collision induced dissociation MS/MS experiments (20 eV, positive ionization mode) of the putative Amadori product FruAcLys isomers with (A) retention time at 6.2 min and (B) 6.6 min, measured with HILIC UHPLC-MS, which were found to be significantly increased in formula-fed infants. (C) Overlaid extracted ion chromatograms of FruAcLys ([M+H]^+^ = 351.1762 ± 0.005 Da) in feces (black) and in a model reaction mixture of L-lysine with glucose, heated for 1 h at 100 °C in water (blue).

**Figure S3.**
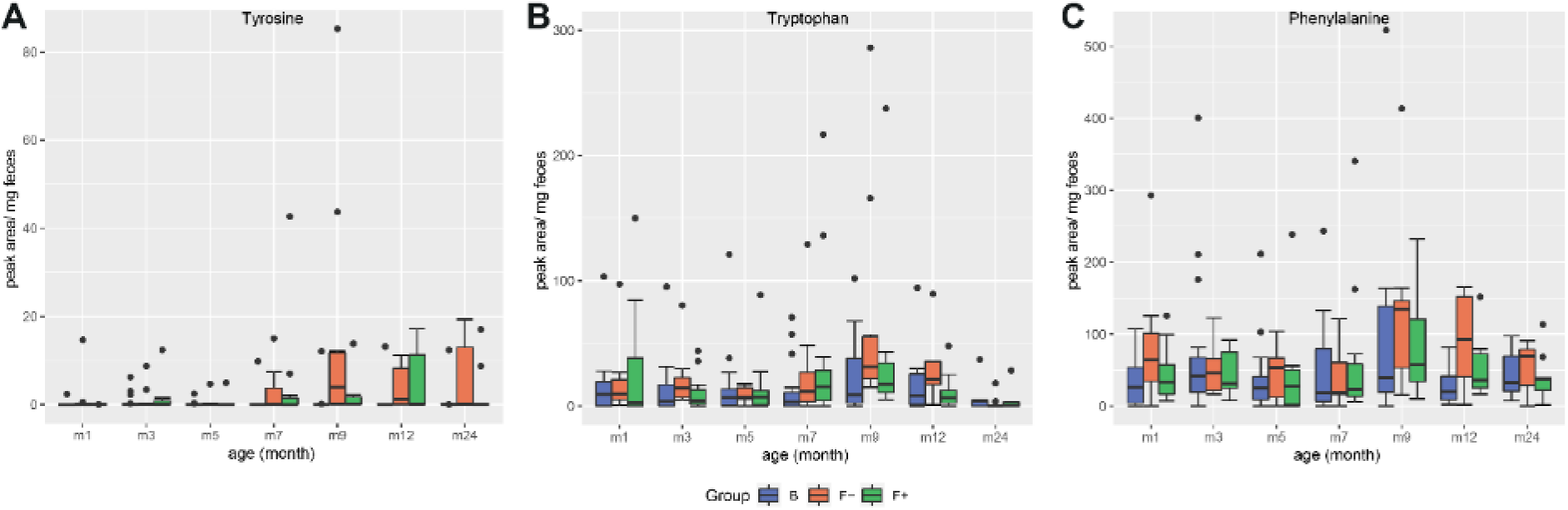
Profiles of the aromatic amino acids (A) tyrosine, (B) tryptophan and (C) phenylalanine in feces of breastfed (group B) and formula-fed infants without (group F-) or with probiotics (group F+) over time. For tryptophan outliers with peak areas > 300 (n = 5) were excluded for visualization.

**Figure S4.**
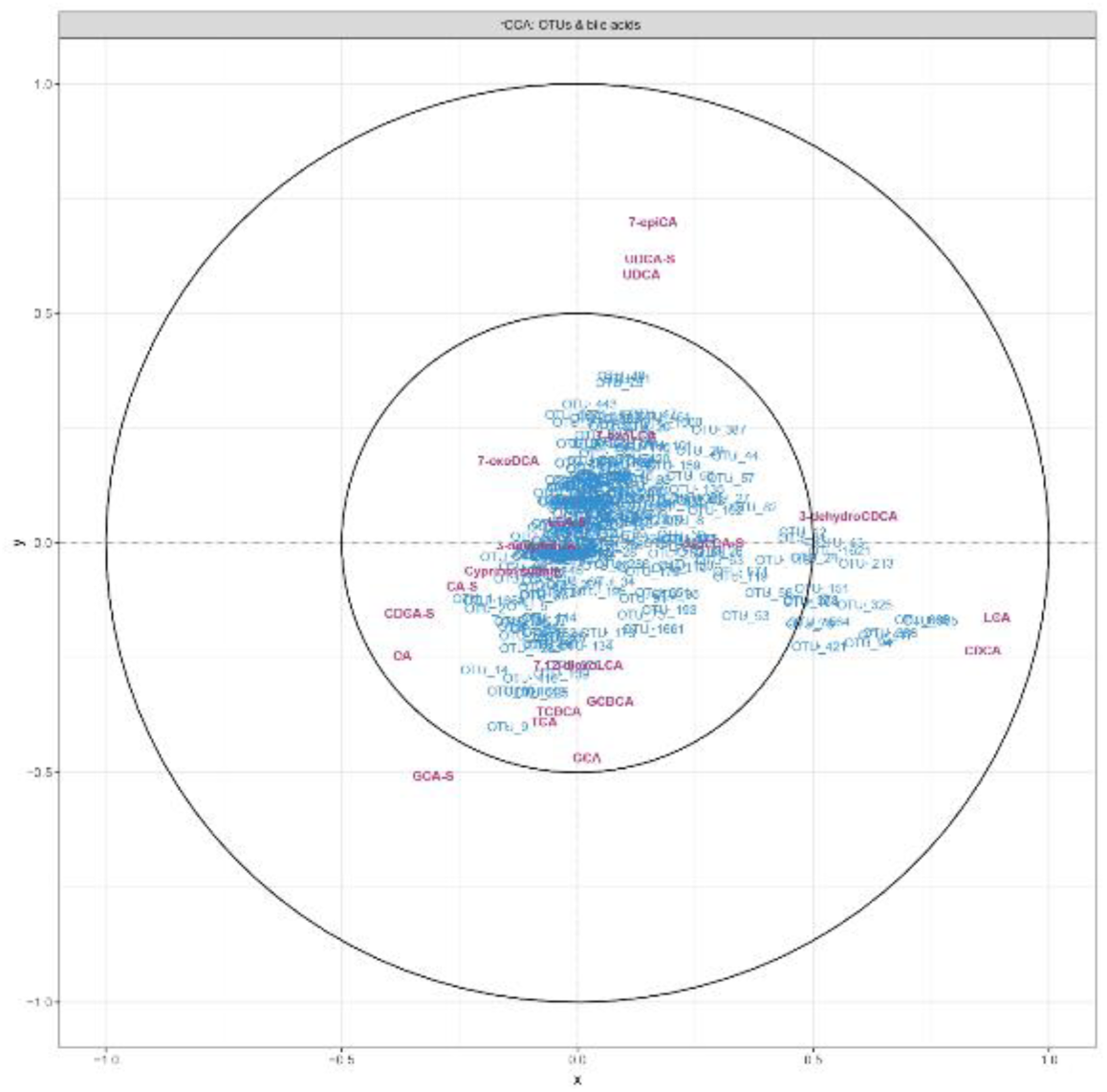
Representation of variables defined by the first two canonical variates from a regularized canonical correlation analysis (rCCA) of bile acids and OTU data from 16S rRNA sequencing detected in feces samples from all infants. Bile acids are colored in purple and OTUs in blue.

**Figure S5.**
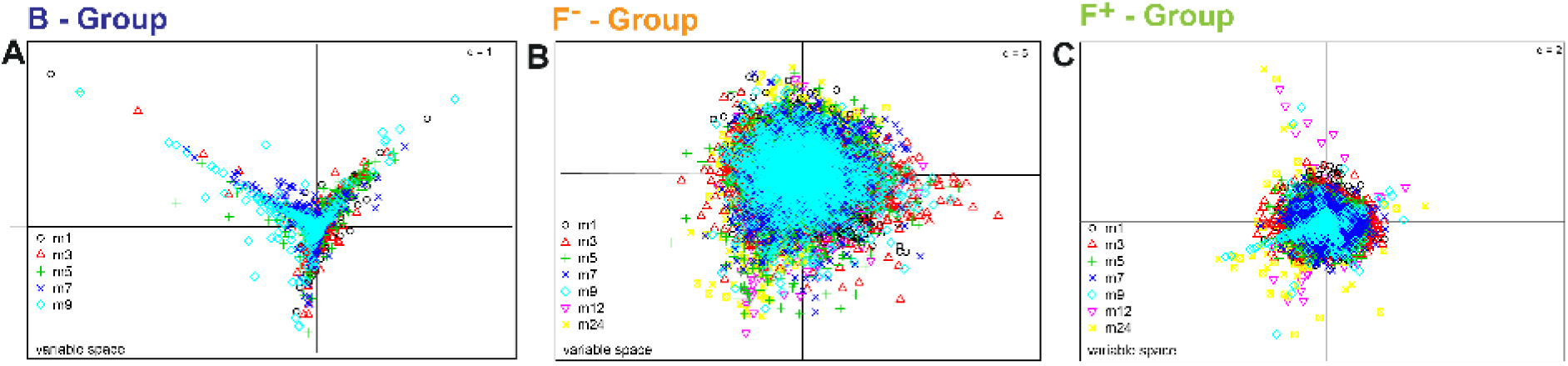
Variable spaces of multiple co-inertia analysis to visualize inter- and intra-individual differences over time, separated for feeding groups; (A) breastfed (B), (B) formula-fed without probiotics (F-) and (C) formula-fed with probiotics (F+). Shapes and colors represent the different time points. For group B only months 1 – 9 were taken into account because of a reduced number of available samples in later months due to weaning.

**Figure S6.**
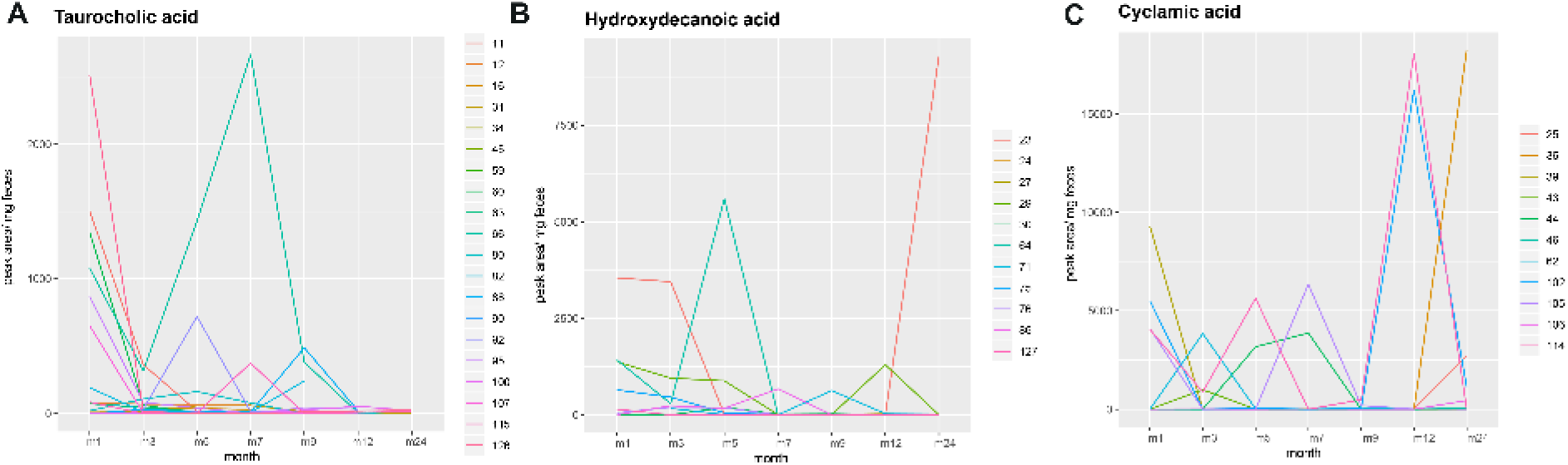
Inter- and intra-individual profiles of three selected metabolites. (A) Increased value of taurocholic acid was found in infant 66 (breastfed) at month 7. (B) High amount of hydroxydecanoic acid in infant 71 (F-) at month 9. (C) High signals of cyclamic acid were detected in infant 25 and 102 (both F+) at month 12.

**Figure S7.**
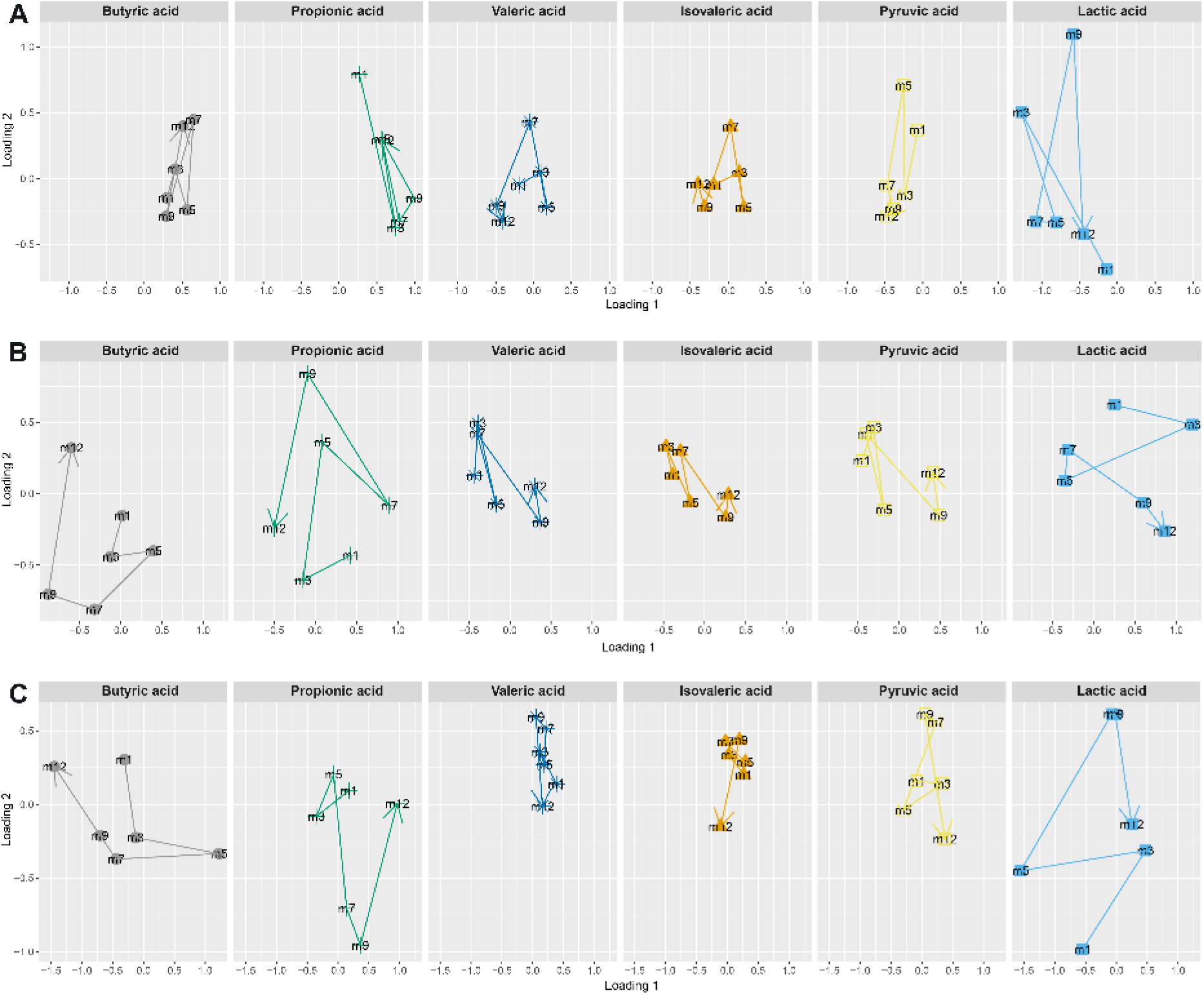
Loading plots of multiple co-inertia analysis of six short chain fatty acids (SCFAs) for breastfed (A), F- (B) and F+ (C), illustrating the dispersion of SCFAs over time. Lactic acid showed high inter- and intra-individual variability in all three groups. Butyric and propionic acid were also highly variable in F- and F+ infants.

## Notes

### Competing Interest Statement

The authors have declared no competing interest.

## REFERENCES

Alnouti, Y. (2009). “Bile Acid Sulfation: A Pathway of Bile Acid Elimination and Detoxification.” Toxicological Sciences 108(2): 225–246.

Aragozzini, F., A. Ferrari, N. Pacini and R. Gualandris (1979). “Indole-3-lactic acid as a tryptophan metabolite produced by Bifidobacterium spp.” Applied and Environmental Microbiology 38(3): 544–546.

Bazanella, M., T. V. Maier, T. Clavel, I. Lagkouvardos, M. Lucio, M. X. Maldonado-Gomez, C. Autran, J. Walter, L. Bode, P. Schmitt-Kopplin and D. Haller (2017). “Randomized controlled trial on the impact of early-life intervention with bifidobacteria on the healthy infant fecal microbiota and metabolome.” Am J Clin Nutr 106(5): 1274–1286.

Begley, M., C. Hill and C. G. M. Gahan (2006). “Bile Salt Hydrolase Activity in Probiotics.” Applied and Environmental Microbiology 72(3): 1729–1738.

Beloborodova, N., I. Bairamov, A. Olenin, V. Shubina, V. Teplova and N. Fedotcheva (2012). “Effect of phenolic acids of microbial origin on production of reactive oxygen species in mitochondria and neutrophils.” Journal of Biomedical Science 19(1): 89.

Beloborodova, N. V., A. S. Khodakova, I. T. Bairamov and A. Y. Olenin (2009). “Microbial origin of phenylcarboxylic acids in the human body.” Biochemistry (Moscow) 74(12): 1350–1355.

Bode, L. (2012). “Human milk oligosaccharides: every baby needs a sugar mama.” Glycobiology 22(9): 1147–1162.

Chow, J., M. R. Panasevich, D. Alexander, B. M. Vester Boler, M. C. Rossoni Serao, T. A. Faber, L. L. Bauer and G. C. Fahey (2014). “Fecal Metabolomics of Healthy Breast-Fed versus Formula-Fed Infants before and during In Vitro Batch Culture Fermentation.” Journal of Proteome Research 13(5): 2534–2542.

Degirolamo, C., S. Rainaldi, F. Bovenga, S. Murzilli and A. Moschetta (2014). “Microbiota modification with probiotics induces hepatic bile acid synthesis via downregulation of the Fxr-Fgf15 axis in mice.” Cell Rep 7(1): 12–18.

Dodd, D., M. H. Spitzer, W. Van Treuren, B. D. Merrill, A. J. Hryckowian, S. K. Higginbottom, A. Le, T. M. Cowan, G. P. Nolan, M. A. Fischbach and J. L. Sonnenburg (2017). “A gut bacterial pathway metabolizes aromatic amino acids into nine circulating metabolites.” Nature 551(7682): 648–652.

Donazzolo, E., A. Gucciardi, D. Mazzier, C. Peggion, P. Pirillo, M. Naturale, A. Moretto and G. Giordano (2017). “Improved synthesis of glycine, taurine and sulfate conjugated bile acids as reference compounds and internal standards for ESI–MS/MS urinary profiling of inborn errors of bile acid synthesis.” Chemistry and Physics of Lipids 204: 43–56.

Eyssen, H. (1973). “Role of the gut microflora in metabolism of lipids and sterols.” Proceedings of the Nutrition Society 32(2): 59–63.

Fiorucci, S. and E. Distrutti (2015). “Bile Acid-Activated Receptors, Intestinal Microbiota, and the Treatment of Metabolic Disorders.” Trends in Molecular Medicine 21(11): 702–714.

Firl, N., H. Kienberger and M. Rychlik (2014). “Validation of the sensitive and accurate quantitation of the fatty acid distribution in bovine milk.” International Dairy Journal 35(2): 139–144.

González, I., K.-A. L. Cao, M. J. Davis and S. Déjean (2012). “Visualising associations between paired ‘omics’ data sets.” BioData mining 5(1): 19–19.

González, I., S. Déjean, P. G. P. Martin and A. Baccini (2008). “CCA: An R Package to Extend Canonical Correlation Analysis.” 2008 23(12): 14.

Grill, J. P., C. Manginot-Dürr, F. Schneider and J. Ballongue (1995). “Bifidobacteria and probiotic effects: Action of Bifidobacterium species on conjugated bile salts.” Current Microbiology 31(1): 23–27.

Hammons, J. L., W. E. Jordan, R. L. Stewart, J. D. Taulbee and R. W. Berg (1988). “Age and Diet Effects on Fecal Bile Acids in Infants.” Journal of Pediatric Gastroenterology and Nutrition 7(1): 30–38.

Hascoët, J.-M., M. Chauvin, C. Pierret, S. Skweres, L.-D. Van Egroo, C. Rougé and P. Franck (2019). “Impact of Maternal Nutrition and Perinatal Factors on Breast Milk Composition after Premature Delivery.” Nutrients 11(2): 366.

Hegele, J., T. Buetler and T. Delatour (2008). “Comparative LC–MS/MS profiling of free and protein-bound early and advanced glycation-induced lysine modifications in dairy products.” Analytica Chimica Acta 617(1): 85–96.

Henning, C., M. Smuda, M. Girndt, C. Ulrich and M. A. Glomb (2011). “Molecular Basis of Maillard Amide-Advanced Glycation End Product (AGE) Formation in Vivo.” Journal of Biological Chemistry 286(52): 44350–44356.

Heubi, J. E., W. F. Balistreri and F. J. Suchy (1982). “Bile salt metabolism in the first year of life.” The Journal of Laboratory and Clinical Medicine 100(1): 127–136.

Hewelt-Belka, W., D. Garwolińska, M. Belka, T. Bączek, J. Namieśnik and A. Kot-Wasik (2019). “A new dilution-enrichment sample preparation strategy for expanded metabolome monitoring of human breast milk that overcomes the simultaneous presence of low- and high-abundance lipid species.” Food Chemistry 288: 154–161.

Huang, C. T., J. T. Rodriguez, W. E. Woodward and B. L. Nichols (1976). “Comparison of patterns of fecal bile acid and neutral sterol between children and adults.” Am J Clin Nutr 29(11): 1196–1203.

John, A., R. Sun, L. Maillart, A. Schaefer, E. Hamilton Spence and M. T. Perrin (2019). “Macronutrient variability in human milk from donors to a milk bank: Implications for feeding preterm infants.” PLOS ONE 14(1): e0210610.

Kim, G.-B., C. M. Miyamoto, E. A. Meighen and B. H. Lee (2004). “Cloning and Characterization of the Bile Salt Hydrolase Genes (*bsh*) from *Bifidobacterium bifidum* Strains.” Applied and Environmental Microbiology 70(9): 5603–5612.

Koletzko, B., S. Baker, G. Cleghorn, U. F. Neto, S. Gopalan, O. Hernell, Q. S. Hock, P. Jirapinyo, B. Lonnerdal, P. Pencharz, H. Pzyrembel, J. Ramirez-Mayans, R. Shamir, D. Turck, Y. Yamashiro and D. Zong-Yi (2005). “Global standard for the composition of infant formula: recommendations of an ESPGHAN coordinated international expert group.” J Pediatr Gastroenterol Nutr 41(5): 584–599.

Lester, R., J. St Pyrek, J. M. Little and E. W. Adcock (1983). “Diversity of bile acids in the fetus and newborn infant.” J Pediatr Gastroenterol Nutr 2(2): 355–364.

Lewis, Z. T., S. M. Totten, J. T. Smilowitz, M. Popovic, E. Parker, D. G. Lemay, M. L. Van Tassell, M. J. Miller, Y.-S. Jin, J. B. German, C. B. Lebrilla and D. A. Mills (2015). “Maternal fucosyltransferase 2 status affects the gut bifidobacterial communities of breastfed infants.” Microbiome 3(1): 13.

Martin, C. R., P.-R. Ling and G. L. Blackburn (2016). “Review of Infant Feeding: Key Features of Breast Milk and Infant Formula.” Nutrients 8(5): 279.

McOrist, A. L., R. B. Miller, A. R. Bird, J. B. Keogh, M. Noakes, D. L. Topping and M. A. Conlon (2011). “Fecal Butyrate Levels Vary Widely among Individuals but Are Usually Increased by a Diet High in Resistant Starch.” The Journal of Nutrition 141(5): 883–889.

Meng, C., B. Kuster, A. C. Culhane and A. M. Gholami (2014). “A multivariate approach to the integration of multi-omics datasets.” BMC Bioinformatics 15: 162.

Meng, C., B. Kuster, A. C. Culhane and A. M. Gholami (2014). “A multivariate approach to the integration of multi-omics datasets.” BMC bioinformatics 15: 162–162.

Neelima, R. Sharma, Y. S. Rajput and B. Mann (2013). “Chemical and functional properties of glycomacropeptide (GMP) and its role in the detection of cheese whey adulteration in milk: a review.” Dairy science & technology 93(1): 21–43.

Pischetsrieder, M. and T. Henle (2012). “Glycation products in infant formulas: chemical, analytical and physiological aspects.” Amino Acids 42(4): 1111–1118.

Poroyko, V., M. Morowitz, T. Bell, A. Ulanov, M. Wang, S. Donovan, N. Bao, S. Gu, L. Hong, J. C. Alverdy, J. Bergelson and D. C. Liu (2011). “Diet creates metabolic niches in the “inmature gut” that shape microbial communities.” Nutricion Hospitalaria 26(6): 1283–1295.

Pranger, I. G., E. Corpeleijn, F. A. J. Muskiet, I. P. Kema, C. Singh-Povel and S. J. L. Bakker (2019). “Circulating fatty acids as biomarkers of dairy fat intake: data from the lifelines biobank and cohort study.” Biomarkers 24(4): 360–372.

Prentice, A. (1996). “Constituents of Human Milk.” Food and Nutrition Bulletin 17(4): 1–10.

Rigo, J., G. Boehm, G. Georgi, J. Jelinek, K. Nyambugabo, G. Sawatzki and F. Studzinski (2001). “An Infant Formula Free of Glycomacropeptide Prevents Hyperthreoninemia in Formula-Fed Preterm Infants.” Journal of Pediatric Gastroenterology and Nutrition 32(2): 127–130.

Robinson, S. M. (2015). “Infant nutrition and lifelong health: current perspectives and future challenges.” J Dev Orig Health Dis 6(5): 384–389.

Ruan, D. W., Hui; Cheng, Faliang (2018). The Maillard Reaction in Food Chemistry: Current Technology and Applications.

Sillner, N., A. Walker, E.-M. Harrieder, P. Schmitt-Kopplin and M. Witting (2019). “Development and application of a HILIC UHPLC-MS method for polar fecal metabolome profiling.” Journal of Chromatography B 1109: 142–148.

Sillner, N., A. Walker, D. Hemmler, M. Bazanella, S. S. Heinzmann, D. Haller and P. Schmitt-Kopplin (2019). “Milk-Derived Amadori Products in Feces of Formula-Fed Infants.” J Agric Food Chem 67(28): 8061–8069.

Sillner, N., A. Walker, W. Koch, M. Witting and P. Schmitt-Kopplin (2018). “Metformin impacts cecal bile acid profiles in mice.” Journal of Chromatography B 1083: 35–43.

Smuda, M., M. Voigt and M. A. Glomb (2010). “Degradation of 1-Deoxy-d-erythro-hexo-2,3-diulose in the Presence of Lysine Leads to Formation of Carboxylic Acid Amides.” Journal of Agricultural and Food Chemistry 58(10): 6458–6464.

Sperisen, P., O. Cominetti and F.-P. Martin (2015). “Longitudinal omics modeling and integration in clinical metabonomics research: challenges in childhood metabolic health research.” Frontiers in Molecular Biosciences 2: 44.

Tanaka, H., H. Hashiba, J. Kok and I. Mierau (2000). “Bile Salt Hydrolase of Bifidobacterium longum — Biochemical and Genetic Characterization.” Applied and Environmental Microbiology 66(6): 2502–2512.

Villaseñor, A., I. Garcia-Perez, A. Garcia, J. M. Posma, M. Fernández-López, A. J. Nicholas, N. Modi, E. Holmes and C. Barbas (2014). “Breast Milk Metabolome Characterization in a Single-Phase Extraction, Multiplatform Analytical Approach.” Analytical Chemistry 86(16): 8245–8252.

Wandro, S., S. Osborne, C. Enriquez, C. Bixby, A. Arrieta and K. Whiteson (2018). “The Microbiome and Metabolome of Preterm Infant Stool Are Personalized and Not Driven by Health Outcomes, Including Necrotizing Enterocolitis and Late-Onset Sepsis.” mSphere 3(3): e00104–00118.

Wang, J., Y.-M. Lu, B.-Z. Liu and H.-Y. He (2008). “Electrospray positive ionization tandem mass spectrometry of Amadori compounds.” Journal of Mass Spectrometry 43(2): 262–264.

WHO (n.d.). “Global Strategy for Infant and Young Child Feeding.”

Wild, J., M. Shanmuganathan, M. Hayashi, M. Potter and P. Britz-McKibbin (2019). “Metabolomics for improved treatment monitoring of phenylketonuria: urinary biomarkers for non-invasive assessment of dietary adherence and nutritional deficiencies.” Analyst 144(22): 6595–6608.

Wishart, D. S., Y. D. Feunang, A. Marcu, A. C. Guo, K. Liang, R. Vazquez-Fresno, T. Sajed, D. Johnson, C. Li, N. Karu, Z. Sayeeda, E. Lo, N. Assempour, M. Berjanskii, S. Singhal, D. Arndt, Y. Liang, H. Badran, J. Grant, A. Serra-Cayuela, Y. Liu, R. Mandal, V. Neveu, A. Pon, C. Knox, M. Wilson, C. Manach and A. Scalbert (2018). “HMDB 4.0: the human metabolome database for 2018.” Nucleic Acids Res 46(D1): D608–d617.

Zhou, W., M. R. Sailani, K. Contrepois, Y. Zhou, S. Ahadi, S. R. Leopold, M. J. Zhang, V. Rao, M. Avina, T. Mishra, J. Johnson, B. Lee-McMullen, S. Chen, A. A. Metwally, T. D. B. Tran, H. Nguyen, X. Zhou, B. Albright, B.-Y. Hong, L. Petersen, E. Bautista, B. Hanson, L. Chen, D. Spakowicz, A. Bahmani, D. Salins, B. Leopold, M. Ashland, O. Dagan-Rosenfeld, S. Rego, P. Limcaoco, E. Colbert, C. Allister, D. Perelman, C. Craig, E. Wei, H. Chaib, D. Hornburg, J. Dunn, L. Liang, S. M. S.-F. Rose, K. Kukurba, B. Piening, H. Rost, D. Tse, T. McLaughlin, E. Sodergren, G. M. Weinstock and M. Snyder (2019). “Longitudinal multi-omics of host–microbe dynamics in prediabetes.” Nature 569(7758): 663–671.

